# A validation scale to determine the readiness of environmental DNA assays for routine species monitoring

**DOI:** 10.1101/2020.04.27.063990

**Authors:** Bettina Thalinger, Kristy Deiner, Lynsey R. Harper, Helen C. Rees, Rosetta C. Blackman, Daniela Sint, Michael Traugott, Caren S. Goldberg, Kat Bruce

**Author notes:** **Corresponding author:** Bettina Thalinger, Centre for Biodiversity Genomics, University of Guelph, 50 Stone Road E, N1G 2W1, Guelph, Ontario, Canada.

## Abstract

The use of environmental DNA (eDNA) analysis for species monitoring requires rigorous validation - from field sampling to the analysis of PCR-based results - for meaningful application and interpretation. Assays targeting eDNA released by individual species are typically validated with no predefined criteria to answer specific research questions in one ecosystem. Hence, the general applicability of assays as well as associated uncertainties and limitations, often remain undetermined. The absence of clear guidelines for assay validation prevents targeted eDNA assays from being incorporated into species monitoring and policy; thus, their establishment is essential for realizing the potential of eDNA-based surveys. We describe the measures and tests necessary for successful validation of targeted eDNA assays and the associated pitfalls to form the basis of guidelines. A list of 122 variables was compiled, consolidated into 14 thematic blocks, (e.g. “*in silico* analysis”), and arranged on a 5-level validation scale from “incomplete” to “operational” with defined minimum validation criteria for each level. These variables were evaluated for 546 published single-species assays. The resulting dataset was used to provide an overview of current validation practices and test the applicability of the validation scale for future assay rating. Of the 122 variables, 20% to 76% were reported; the majority (30%) of investigated assays were classified as Level 1 (incomplete), and 15% did not achieve this first level. These assays were characterised by minimal *in silico* and *in vitro* testing, but their share in annually published eDNA assays has declined since 2014. The meta-analysis demonstrates the suitability of the 5-level validation scale for assessing targeted eDNA assays. It is a user-friendly tool to evaluate previously published assays for future research and routine monitoring, while also enabling the appropriate interpretation of results. Finally, it provides guidance on validation and reporting standards for newly developed assays.

## 1. Introduction

Determining the occurrence of species is essential for ecology and requires sensitive and accurate detection methods. Within the last decade, species detection from environmental DNA (eDNA; i.e. detection of extra-organismal DNA released by organisms into their environment) has shown great potential for routine species surveys (Rees, Maddison, Middleditch, Patmore, & Gough, 2014; Goldberg et al., 2016; Deiner et al., 2017; Langlois, Allison, Bergman, To, & Helbing, 2020; Sepulveda, Nelson, Jerde, & Luikart, 2020). The interest in molecular species detection has fuelled the development of over 500 assays, reviewed herein, that utilise PCR to amplify DNA or RNA extracted from environmental samples. Generally, “targeted” eDNA assays must be specific to the species of interest and possess high sensitivity to allow detection at low densities, low DNA concentrations, and across spatiotemporal scales (Goldberg et al., 2016; MacDonald & Sarre, 2017).

A targeted eDNA assay encompasses the entire workflow used to detect a species’ DNA from an environmental sample, inclusive of field sampling through to the interpretation of PCR-based results; it does not just consist of the primers and probes. Thus, adherence to workflows will determine the success or failure of an eDNA assay because methodological choices influence performance and sensitivity (e.g. Doi, Takahara, et al., 2015; Tsuji, Takahara, Doi, Shibata, & Yamanaka, 2019). In practice, assays are often validated within a specific system to answer a set question about the target species. Hence, applications beyond this initial development are hampered by the poor understanding of remaining uncertainties, such as the potential for false positives resulting from non-target amplification or contamination, or false negatives resulting from low sensitivity, sample degradation, low DNA yield protocols or inhibition (Goldberg et al., 2016; Lacoursière-Roussel & Deiner, 2019). In a management context, false positives and false negatives may lead to misuse of resources (e.g. funds and personnel) for issues such as rare species protection and invasive species control. Both scenarios foster inaccurate interpretation of results, fuelling arguments against the routine use of eDNA detection for species monitoring (Jerde, 2019).

Aside from a few well-validated eDNA assays already incorporated into routine monitoring, the application of published assays is a minefield for end-users to navigate. We illustrate this through two examples. The assay for great crested newt (*Triturus cristatus* (Laurenti, 1768), a legally protected species in the UK [Natural England, 2015]) was one of the first eDNA assays validated in both laboratory and field trials against conventional tools, demonstrating its potential for routine monitoring (Thomsen et al., 2012; Rees, Bishop, et al., 2014). After successful validation, a national eDNA-based citizen science monitoring scheme was tested and showed that large-scale eDNA sampling can enable distribution modelling (Biggs et al., 2015). These initial studies paved the way for eDNA-based *T. cristatus* detection to inform new policies aimed at providing landscape-level species protection (L. R. Harper, Buxton, et al., 2019). Studies have since investigated optimal methods of eDNA capture, relative abundance and detection probability estimation, and the influence of seasonality as well as biotic and abiotic factors on *T. cristatus* eDNA detection and quantification (Buxton, Groombridge, & Griffiths, 2017; Buxton, Groombridge, Zakaria, & Griffiths, 2017; Buxton, Groombridge, & Griffiths, 2018a, 2018b). Due to these combined efforts, the assay has undergone exemplary validation and has been operational for management since 2014.

Conversely, no assays have been successfully applied to routine monitoring of invasive American crayfish. For example, there is a lack of consensus on a single assay for the signal crayfish (*Pacifastacus leniusculus* (Dana, 1852)). Larson et al. (2017) developed and tested an assay against conventional trapping, but five other assays have also been proposed with differing degrees of validation and divergent *in silico* and *in vitro* approaches (Agersnap et al., 2017; Dunn, Priestley, Herraiz, Arnold, & Savolainen, 2017; K. J. Harper, Anucha, Turnbull, Bean, & Leaver, 2018; Mauvisseau et al., 2018; Robinson, Uren Webster, Cable, James, & Consuegra, 2018). The *P. leniusculus* assays were developed using a variety of strategies for eDNA sampling, capture, extraction, qPCR of different genetic markers, and applied across genetically diverse populations within the species range. Due to this substantial methodological variability, direct comparisons between results obtained from these assays are impossible. Therefore, the *P. leniusculus* assays represent a minefield for end-users, despite the need for accurate and sensitive tools to enable actionable species management of this invasive species in Europe, Japan, and California (USA).

These case studies exemplify how consensus and dissent in assay validation can influence the implementation of eDNA analysis for species monitoring. Developing guidelines to determine the suitability of eDNA assays for end-users will therefore ensure that ecological insights or management decisions are based on robust molecular analyses with quantifiable uncertainties and clear inference limits (Goldberg et al., 2016; MacDonald & Sarre, 2017; Nicholson et al., 2020; Sepulveda et al., 2020). Here, we describe the general validation process for targeted PCR-based methods and examine the extent of assay validation and reporting in the eDNA literature. We present an eDNA assay validation scale, which establishes criteria to enable the classification of assays based on their accuracy and sensitivity for single-species detection. To demonstrate the utility of the scale, we performed a meta-analysis of targeted eDNA assays published in 327 papers as of 11 April 2019 (546 assays). By placing an eDNA assay on the validation scale, end-users can determine the recommended scenarios for application and improve assay performance with further validation.

## 2. Criteria and principles of validation

### 2.1. General requirements for an eDNA laboratory

All laboratory activities are subject to error. In order to have confidence in results, quality standards and good practices are required in diagnostic laboratory environments (e.g. World Health Organization, 2011; Halling, Schrijver, & Persons, 2012). Although few eDNA-processing laboratories will employ ‘ancient DNA’ practices (e.g. full body suit and positive air pressure with HEPA filtered inflow), all laboratories conducting eDNA analysis should utilise a unidirectional workflow where pre-PCR steps are performed in separate laboratories dedicated to low DNA quality and quantity (Goldberg et al., 2016). Completely standardised laboratory environments are rare and the use of proficiency tests (as conducted by UK laboratories participating in *T. cristatus* monitoring) can help end-users understand the quality of results obtained among different laboratories. Even results obtained from an extensively validated assay can be questionable when they are not produced within a suitable laboratory environment (Goldberg et al., 2016).

### 2.2. Reporting standards for in silico, in vitro and in situ validation of assays

Targeted eDNA assay validation is a multi-step process. It can be divided into *in silico* validation (i.e. computer-based tests for primer specificity), *in vitro* validation (i.e. laboratory tests with reference tissue samples) and *in situ* validation (i.e. field tests with eDNA samples) (see Goldberg et al., 2016; MacDonald & Sarre, 2017; Langlois et al., 2020; So, Fong, Lam, & Dudgeon, 2020). Understanding the utility of an assay requires both knowledge of the context in which it has been designed, and a broader understanding of how it was developed. Here, we give a brief overview of what key steps comprise assay validation, with a focus on troubleshooting steps that may be necessary when applying previously published assays to new locations or with modified laboratory practices (Table 1).

**Table 1:**
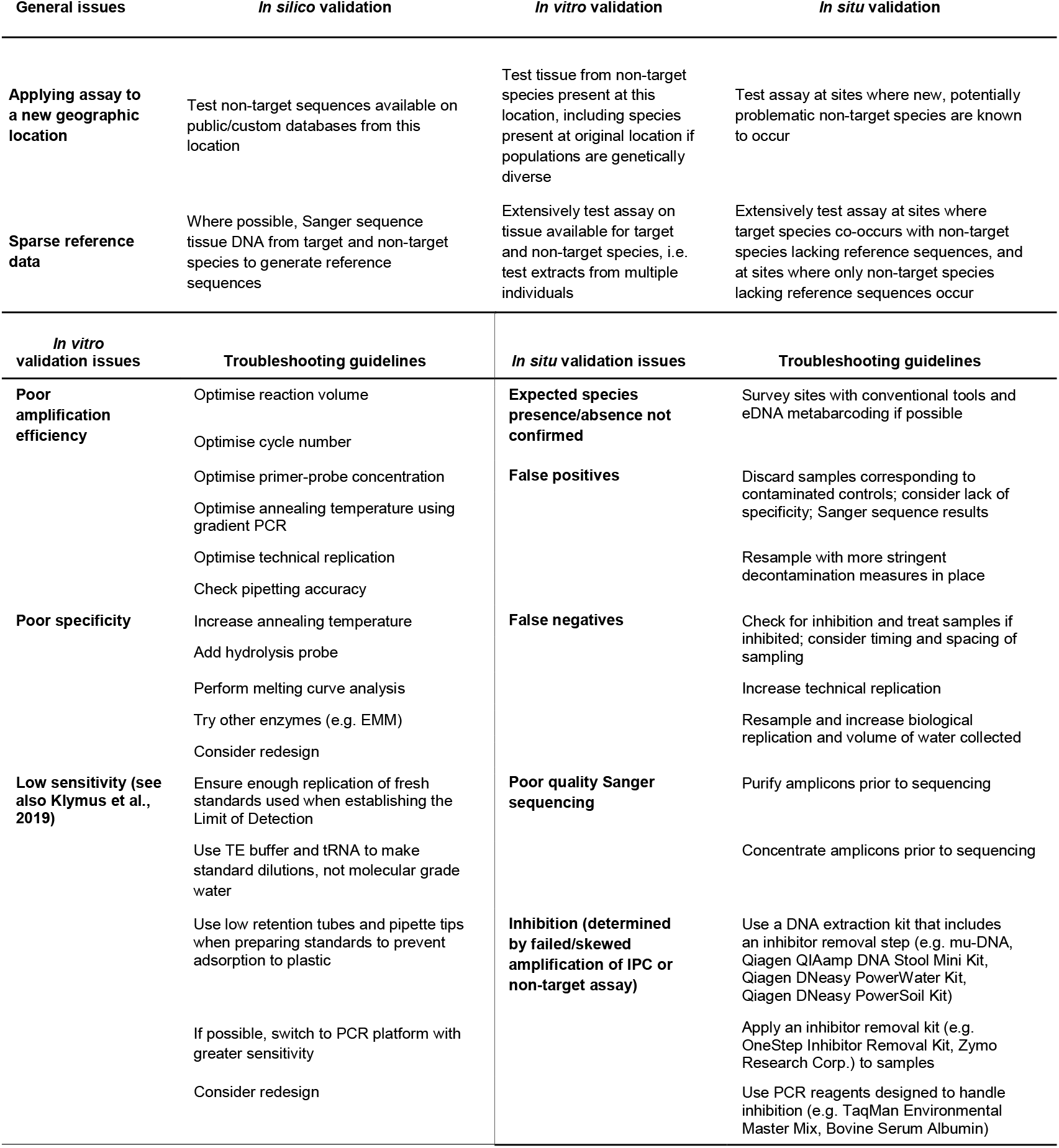
A guideline for troubleshooting at different stages of the validation process.

The first step is *in silico* assay validation, the goal of which is to determine assay specificity based on known sequence diversity. Sequence diversity has three categories, including sequences from: (i) closely related and co-occurring species, (ii) closely related but geographically distinct species, and (iii) distantly related but co-occurring species (i.e. sequences that could co-amplify and produce false positive results for a target species). By checking primer specificity from available sequences, the geographic area of applicability for an assay can be maximised through identifying and removing potential issues of co-amplification. Typically, public or custom databases are used for performing *in silico* amplification (e.g. ecoPCR [Boyer et al., 2016]; primerBlast [Ye et al., 2012]; PrimerTREE [Cannon et al., 2016]; PrimerMiner [Elbrecht & Leese, 2017]). While reference sequence libraries are often far from complete and many of the factors influencing successful PCR amplification cannot be simulated, *in silico* testing provides a first impression of primer performance and should be conducted.

The essential components of *in vitro* assay validation are optimisation, specificity, and sensitivity. Tests with varying PCR chemistry, reaction volume, primer/probe concentration, cycling conditions and technical replication will ensure optimal, standardised and error-free target DNA amplification (Bustin et al., 2009; Wilcox, Carim, McKelvey, Young, & Schwartz, 2015; Goldberg et al., 2016). The assay must then be tested against closely related and co-occurring non-target taxa to ensure specificity, which is not automatically guaranteed after successful *in silico* testing (Goldberg et al., 2016; So et al., 2020). Ideally, tissue-derived DNA samples from multiple individuals spanning a defined geographic area are tested to ensure the assay is robust to genetic variants of target and non-target species. Amplicons should be Sanger sequenced to confirm species identity (Goldberg et al., 2016), although short fragments (<100 bp) have limited sequence length available for species determination (Meusnier et al., 2008). Next, the Limit of Detection (LOD) must be determined to assess assay sensitivity, and the Limit of Quantification (LOQ) determined, if the measurement of eDNA quantity is desired. Generally, these values are obtained using a dilution series of quantified DNA amplicons or synthesized gene fragments (e.g. IDT gBlocks™ Gene Fragment) based on public or *de novo* reference sequences (Langlois et al., 2020). The LOD and LOQ have various definitions in the eDNA literature, but were recently standardised by Klymus et al. (2019), where LOD is the lowest standard concentration at which 95% of technical replicates amplify and LOQ is the lowest standard concentration for which the coefficient of variation (CV) value is <35%. Unfortunately, the existence of past definitions requires the final LOD and LOQ to be reported as well as the definition used. We note that these metrics apply directly to the assay as developed and assume no interference during PCR from the rest of the species’ genome (i.e. if a gBlock is used), other genetic material, or inhibitory compounds.

Finally, the assay must be validated *in situ* by surveying sites with and without the target species (Goldberg et al., 2016). It must be tested against conventional tools for presence/absence detection and tests for estimation of relative abundance/biomass are advisable. Assays are deemed successful if eDNA and conventional detections concur at occupied sites and no eDNA detections are observed at definitively unoccupied sites. Sanger sequencing of eDNA amplicons can provide additional evidence but cannot distinguish sample contamination from true detections (Goldberg et al., 2016). Besides screening for the target species, negative eDNA samples (or all eDNA samples if quantification is necessary) should be tested for inhibition. This requires an Internal Positive Control (IPC) assay for synthetic DNA (e.g. ThermoFisher) or an assay for non-target species using exogenous or endogenous DNA (Goldberg et al., 2016; Veldhoen et al., 2016; Doi et al., 2017; Furlan & Gleeson, 2017).

Advanced *in situ* validation may investigate the influence of biotic (e.g. abundance, biomass, life stages, microbial activity) and abiotic (e.g. temperature, pH, ultraviolet light, salinity) factors influencing eDNA origin, state, fate, and transport (Barnes & Turner, 2016; Lacoursière-Roussel & Deiner, 2019; Wang et al., 2021). Assays that account for spatial (e.g. shoreline versus offshore) and temporal (e.g. summer versus winter) variation in eDNA distribution and abundance due to the ecology of a species can be implemented with greater confidence (de Souza, Godwin, Renshaw, & Larson, 2016; Lawson Handley et al., 2019). Occupancy modelling using eDNA data is desirable as it accounts for detection probability while estimating site occupancy, even if all field samples from a site return negative. Hierarchical models that incorporate eDNA occupancy and detection probabilities at site, sample, and technical replicate levels are most accurate and can be implemented in software such as R (e.g. package “eDNAoccupancy” [Dorazio & Erickson, 2018]) or PRESENCE (MacKenzie et al., 2002). However, model assumptions regarding false positives should be carefully considered.

## 3. Types and trade-offs of targeted eDNA detection methods

Amid the processing chain (i.e. sampling to data analysis) for a targeted eDNA assay, PCR warrants extra consideration as the technological spectrum and potential for variation is enormous. Previous publications have typically defined an assay as the primers (and probe) required for DNA amplification and associated visualisation (e.g. agarose gel electrophoresis, qPCR instrumentation). However, differences between the multiple detection instruments and chemistry used in combination with species-specific primer (and probe) sets can fundamentally change the sensitivity and specificity of targeted eDNA assays. Table 2 provides an overview of amplification types and their associated trade-offs.

**Table 2:** Detection methods and their trade-offs used for targeted eDNA assays. Abbreviations are as follows: polymerase chain reaction (PCR), capillary electrophoresis (CE), quantitative (q), digital (d), intercalating dye (ID), loop-mediated isothermal amplification (LAMP).

| Trade-offs | Endpoint PCR-gel | Endpoint PCR-CE | qPCR-ID | qPCR-probe | dPCR-ID | dPCR-probe | LAMP |
| --- | --- | --- | --- | --- | --- | --- | --- |
| **Quantification** | no | yes<br>(limited precision) | yes | yes | yes | yes | yes, with real-time monitoring |
| **LOD (sensitivity)** | medium - high DNA concentration<br>(depending on which intercalating dye) | low - medium DNA concentration (with a well-designed assay) | low DNA quantities | low DNA quantities | absolute DNA quantities | absolute DNA quantities | medium DNA concentration |
| **Specificity** | medium | medium | medium | high | medium | high | high |
| **Cost reagents** | low | medium | high | high | high | very high | medium |
| **Cost equipment** | low | medium | medium | medium | high | high | low - medium<br>(depending on LAMP cyclers) |
| **Multiplex possible** | yes, if different sizes are used | yes, if different sizes or fluorophores are used | difficult | yes, if different fluorophores are used | difficult | yes, if different fluorophores are used | yes |
| **Time (PCR to data)** | slow | slow | medium | medium | slow | slow | fast (15-30 min) |
| **Assay type transferable** | yes | yes | yes | yes | yes, to qPCR | yes, to qPCR | not easily |
| **Effort to design assay** | low - medium | low - medium | high | high | high | high | high |
| **Specialised software** | advisable | advisable | advisable | required | advisable | required | required |
| **Sequencing confirmation** | yes | yes | yes | yes | limited | limited | limited |

Many assays - especially those published in earlier years - use endpoint PCR. However, most assays to date employ real-time quantitative (q)PCR allowing for greater sensitivity and quantitative data. More recent publications have used digital (d)PCR for absolute quantification. Alternatively, LAMP and CRISPR have been shown to be suitable for eDNA applications, decreasing the requirements of in-field testing equipment (M. R. Williams et al., 2017; M. A. Williams et al., 2019). A few publications use alternative methods such as PCR combined with restriction fragment length polymorphism (RFLP). All amplification types (Table 2) enable distribution and occupancy modelling, provided enough biological and technical replication is employed (Hunter et al., 2015; Goldberg, Strickler, & Fremier, 2018; Wilcox et al., 2018). However, endpoint PCR in combination with agarose gels is the only type where a detection limit cannot be set objectively (low sensitivity) and which does not provide estimates of DNA copy number (Doi, Uchii, et al., 2015; Yamanaka & Minamoto, 2016; Hunter et al., 2017; Thalinger, Wolf, Traugott, & Wanzenböck, 2019). Depending on the management context and study size, the optimal detection instrument can vary, albeit technological advances and required accuracy will continue to shift the focus away from endpoint PCR.

## 4. Evaluating the current status of assay validation

To understand current validation practices for targeted eDNA assays, we generated an extensive list of variables deemed important for assay validation by 35 experts in the field of targeted eDNA detection whom convened at a DNAqua-Net EU COST Action (Leese et al., 2016) workshop held on 26-27 March 2018 at the University of Innsbruck, Austria. The variables in this list consist of 18 categorical variables (e.g. species identity, target gene, sample type) and 104 binomial variables directly associated with the eDNA processing chain, from primer design to interpretation of field study results (SI1).

A comprehensive literature database for targeted eDNA assays was built in three steps. First, we included all papers listed on the ‘eDNA assays’ web page (https://labs.wsu.edu/edna/edna-assays/) as of 10 April 2019. Second, we conducted a Web of Science literature search on 11 April 2019, including the search terms “environmental DNA” and “eDNA” but excluding terms associated with microbial organisms and metabarcoding (SI2A). Third, the resulting 660 Web of Science entries were manually checked for suitability (i.e. macrobial target organism, targeted eDNA detection intended) leading to a combined database of 327 papers. For each of the assays contained in these papers, the 122 variables were recorded in a checklist by one of the authors. Before data entry, all authors validated the same four papers (Deiner & Altermatt, 2014; Rees, Bishop, et al., 2014; Thalinger et al., 2016; L. R. Harper, Griffiths, et al., 2019) to ensure recorder standardisation. Validation efforts were classified as 1 for “tests done, or parameter reported”, 0 for “variable not reported, or no testing done”, and NA in cases where the respective variable did not match the assay type (e.g. filter type when samples were precipitated). When an assay was used in multiple papers, all validation efforts were summarised in one entry and the literature database was extended with the papers reporting primer sequences or other methodological aspects. As the type of amplification is important for assay validation, primer pairs used on multiple detection platforms were given separate database entries per amplification type. However, because most assays developed were presented in one publication, we did not account for slight variations in other aspects of the workflow (e.g. different extraction method, different filter type). After recording the values for each eDNA assay using the validation checklist (SI1), each author scored the assay intuitively based on a preliminary version of the validation scale (see section 5). The resulting database of 122 variables for each assay was the basis for all further analyses using R (R Core Team, 2020) and associated packages (SI2B).

Altogether, 546 assays from 327 papers were assessed. Of these assays, 227 were designed to detect fish species and 74 were designed to detect amphibian species; hence, it is unsurprising that ~80% of assays utilised water sampling. Fourteen percent of the assays were tested on tissue only and few assays were optimised for other sample types such as aerosol, sediment, snow or soil. More than 80% of assays were reported in only one paper, and most were designed for qPCR (~60%) or endpoint PCR (~35%) platforms. The cytochrome c oxidase subunit I (*COI*) gene was the most popular (>40%) genetic marker, followed by the cytochrome b (*cytb*) gene (~23%) (SI3).

## 5. The 5-level assay validation scheme

To enable standardised assay validation and reporting in the future, we assigned the assay-specific variables to 14 thematic blocks such as “*in silico* analysis”, “PCR”, or “extensive field testing of environmental samples” (Table 3, Fig. 1). Each of the variable blocks contains between three and 26 variables (SI1). Some of the variable blocks summarize basic practices (e.g. “*in silico* analysis”) while others describe advanced assay validation (e.g. “detection probability estimation from statistical modelling”). To simplify the reporting of an assay’s validation level in the future, the blocks were placed on a five-level scale enabling the categorisation of assays from Level 1 (incomplete) to Level 5 (operational), and the interpretation of associated field study results (Fig. 1). The scale is additive, which means for an assay to be placed at Level 3, it must fulfil the reporting requirements of Levels 1, 2, and 3. Level 1 (incomplete) summarizes assays for which basic *in silico* analysis, target tissue testing and general information regarding PCR were reported. Level 2 (partial) – assays were characterized by comprehensive reporting of PCR conditions and *in vitro* testing on closely related non-target species. For assays placed at Level 3 (essential), the target organism was successfully detected from an environmental sample and the specifics of DNA extraction and concentration of eDNA from the environmental sample (i.e. filtration, precipitation) were reported. The LOD, extensive field testing and *in vitro* testing on co-occurring non-target species were the variable blocks specifically associated with Level 4 (substantial). At Level 5 (operational), the assay has been subjected to comprehensive specificity testing, detection probability estimates from statistical modelling and investigations of ecological and physical factors potentially influencing eDNA in the environment (Table 3, Fig. 1).

**Table 3:** The thematic variable blocks of the 5-level validation scale and their respective minimum criteria.

| Validation level | Variable blocks | Minimum criteria |
| --- | --- | --- |
| Level 1 | *in silico* analysis | target species |
|  | target tissue testing | target tissue |
|  | target tissue PCR | primer (and probe) sequence |
| Level 2 | comprehensive reporting of PCR conditions | DNA extract volume in PCR |
|  | *in vitro* testing on closely related non-target species | any *in vitro* non-target testing |
| Level 3 | extraction method performed on eDNA samples | method of extraction |
|  | concentration of eDNA from environmental sample | filter type or precipitation chemicals |
|  | detection obtained from environmental samples | detection from an environmental sample (artificial or natural habitat) |
| Level 4 | Limit of Detection (LOD) | LOD determined |
|  | extensive field testing of environmental samples | multiple locations or multiple samples |
|  | *in vitro* testing on co-occurring non-target species | any advanced *in vitro* testing |
| Level 5 | comprehensive specificity testing | non-co-occurring/closely related species checked from *in silico* |
|  | detection probability estimation from statistical modelling | any effort made towards detection probability estimation |
|  | understanding ecological and physical factors influencing eDNA in the environment | any factor influencing eDNA in the environment tested |

**Figure 1:**
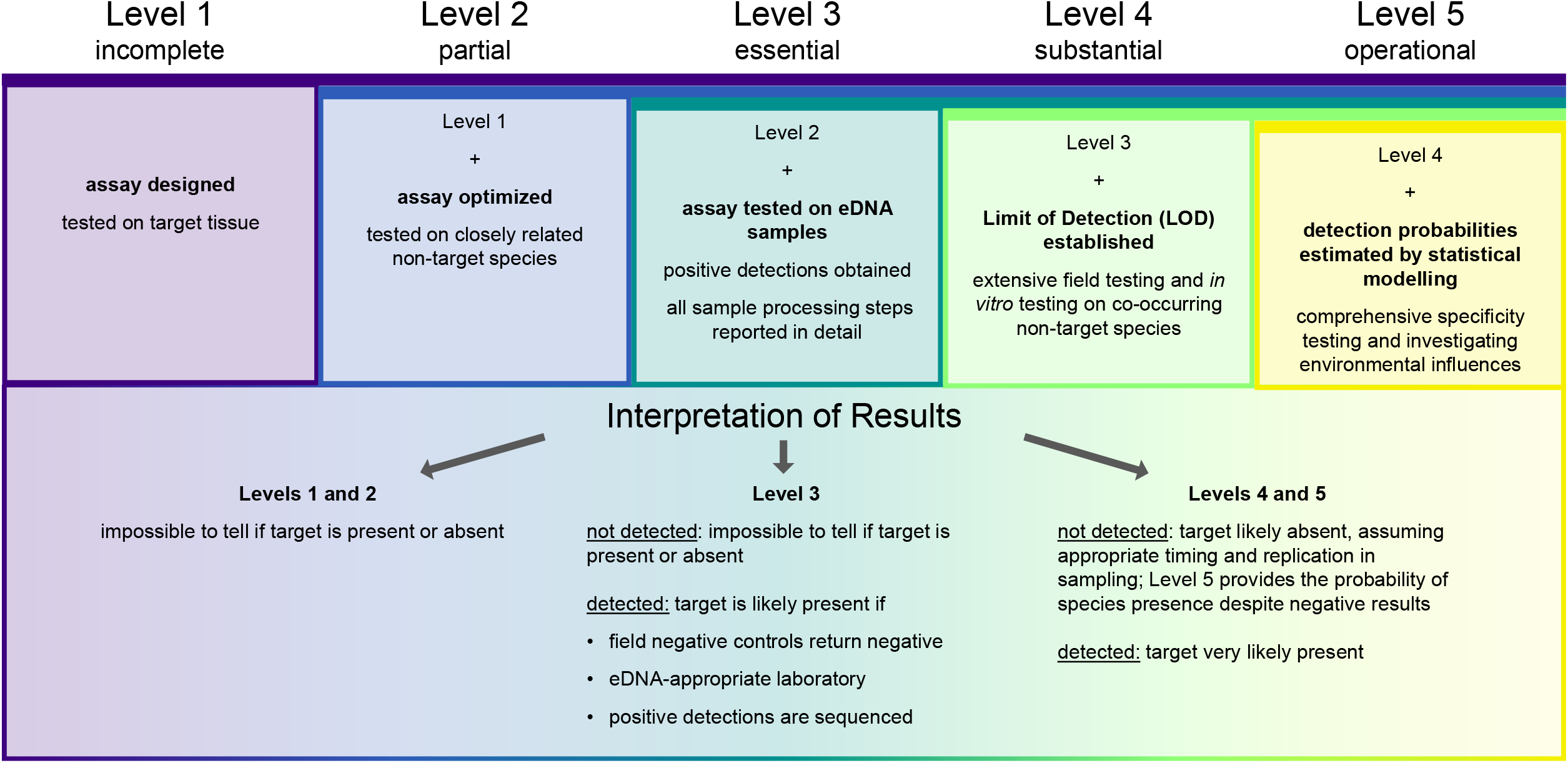
An overview of the 5-level validation scale. For each of the levels (incomplete to operational), the main accomplishments in the validation process and an appropriate interpretation of results are provided.

The placement of assays on this 5-level scale is not straightforward, since each of the 14 variable blocks contains variables associated with either rudimentary or substantial validation and reporting. For instance, the thematic block “concentration of eDNA from environmental sample” contains the variable “volume/weight of environmental sample”, which was reported for almost all assays, but also contains “pressure used for filtration”, which was rarely reported and/or measured. Therefore, a minimum criterion was introduced for each variable block functioning as proof of validation and ensuring standardized placement of assays on the validation scale. For example, “detection from an environmental sample” was used as evidence that some validation had been undertaken in the block “detection obtained from environmental samples” (Table 3). To reach a level on the validation scale, an assay must fulfil all minimum reporting criteria for that level and any preceding levels.

Based on this classification, the results (detected vs. not detected) obtained from eDNA assays become directly interpretable: when Level 1 or Level 2 assays are applied to environmental samples without any further validation steps, it is impossible to tell whether the target species is present or absent independent of the PCR result. Amplifications with a Level 3 assay can be interpreted as “target is likely present”; however, non-amplifications are inconclusive. When Level 4 and Level 5 assays do not lead to amplification, the target is likely absent. Positive PCR results at Level 4 and 5 mean that the target species is almost certainly present (Fig. 1).

For a quantitative analysis of reporting practices, we calculated a total scoring percentage and a block scoring percentage for each assay. The total scoring percentage was defined as the proportion of the 104 binary variables which were reported. For each of the 14 blocks, the block scoring percentage was calculated by dividing the number of variables tested/reported by the complete set of variables associated with the block. Both calculations included only variables relevant to the applied methods (e.g. for assays using filtration, precipitation variables were omitted; see Box 1 for example assays).

Of the 546 assays analysed, the majority (30%) were classed as Level 1. Of the remainder, 15% (*N* = 83) did not fulfil the minimum criteria necessary to reach Level 1, and no assay reached Level 5 (Fig. 1 and Fig. 2A). Newer assays published after 2016 were more likely to reach Level 4, and the percentage of assays failing to reach Level 1 gradually declined since 2014 (Fig. 2B). Generally, the total scoring percentage for all variables in a level increased from Level 0 to Level 4, but its variation was non-uniform with most outliers observed at Level 2 (Fig. 2C). Assays not reaching Level 1 exhibited scoring percentages between ~20% and ~55%, clearly showing the difference between incomplete and partially validated assays, which did not achieve higher levels due to one or several missing validation step(s). Generally, variables associated with lower levels on the validation scale were more likely to be reported or tested. Nevertheless, some variables (e.g. “haplotypes of target tissue” or “pressure used for filtration”), were addressed by fewer than 10% of assays (Fig. 2D).

**Figure 2:**
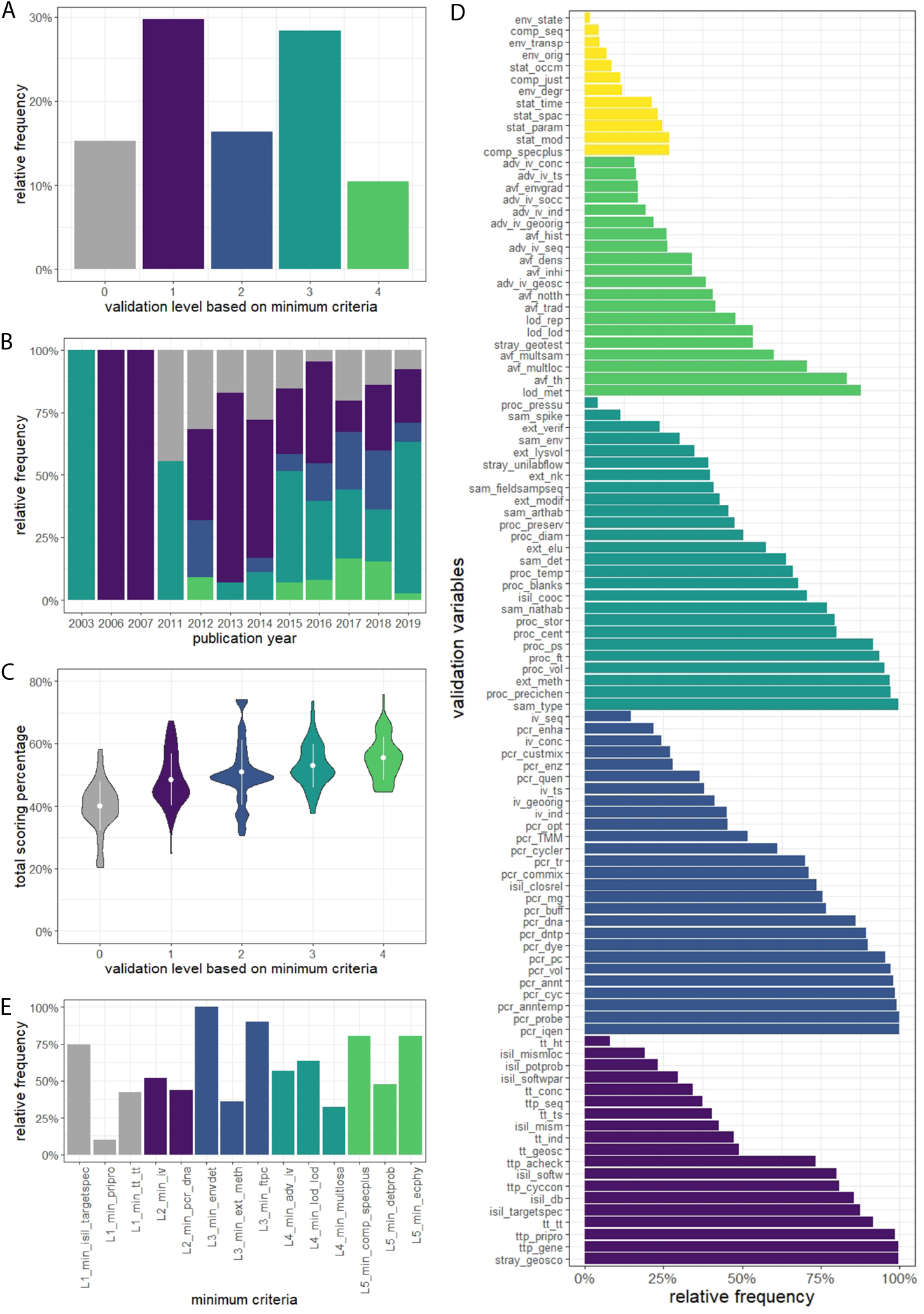
The main outcomes of the meta-analysis based on 546 assays from 327 publications are presented in panels A to E. Assay classification is based on the minimum criteria presented in Figure 1. Level 0 codes for assays that did not reach Level 1 on the validation scale. The colour coding is consistent for all panels: Level 0 (grey), Level 1 (dark purple), Level 2 (blue), Level 3 (turquoise), Level 4 (green), and Level 5 (yellow). Panel A shows the distribution of assays across levels of the validation scale. Panel B displays the percentage of assays (*N* = 546) rated Level 0 to 4 that have been published each year since 2003. Panel C summarises variable reporting per assay level. Panel D shows the percentage of assays reporting a specific variable (colour-coded according to level). Panel E shows the minimum criteria necessary to reach each level of the validation scale, and the percentage of Level 0 to 4 assays that did not report these. All variable abbreviations are listed in SI1.

The rigour of the minimum criteria was evaluated by tallying the not achieved/reported cases. Specifically, 62 assays did not reach Level 1 because the targeted species sequence(s) used in primer design were not reported, which is a prerequisite for this level. Eight assays did not report the primer sequence. The lack of target detection from an environmental sample and omission of filter type or precipitation chemicals were the most restrictive criteria and were not fulfilled by 80 and 43 assays respectively, most of which were exclusively used for tissue tests (Fig. 2E). For assays ranked at Levels 2 and 3, there was agreement between the intuitive assay rating provided by the recorder and that assigned by the objective criteria. For assays placed at Level 1 following objective criteria, authors tended to be more liberal and rated them one or two levels higher (SI4). Finally, a classification tree analysis (De’Ath & Fabricius, 2000) was carried out to identify common characteristics of assays placed at each level of the validation scale (SI5). Most assays failing to reach Level 1 showed distinctly low levels of *in silico* validation. This was also true for most Level 1 assays (*N* = 103), albeit these showed higher levels of target tissue validation (Fig. 3). On the other end of the spectrum, Level 4 assays were characterised by substantial testing or reporting for *in vitro* testing, field sample processing, LOD determination, PCR, and field testing.

**Figure 3:**
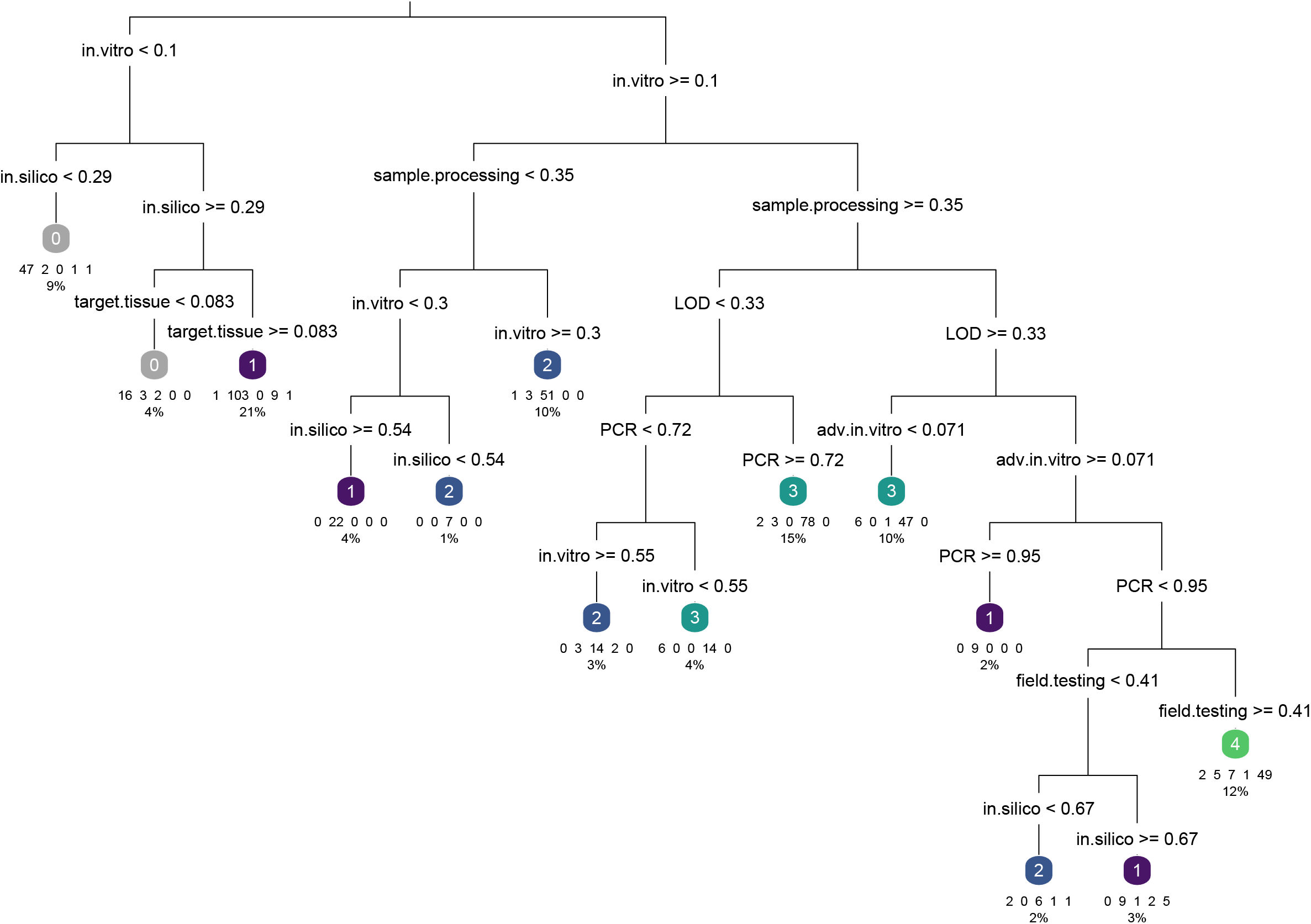
Classification tree analysis identifying the criteria distinguishing assays at different levels of the validation scale. The conditions along the branches show the criteria on which the dataset is split. Numbers in coloured leaves show the validation level of the assays in the respective leaf. Numbers below the leaves represent the number of assays per validation level, summarised inside an individual leaf. The displayed percentage is the proportion of assays summarised per leaf.

## 6. Conclusions and outlook

The validation scale and reporting standards developed here are an inclusive set of guidelines for targeted eDNA assays. They can facilitate communication between the scientific community, commercial providers and government agencies, and provide guidelines regarding the application of previously published assays or the development and publication of new assays (Sepulveda et al., 2020). One needs to acknowledge that a strict standardisation of eDNA assays will not be possible due to applications for manifold taxa in diverse ecosystems combined with technological advances. By checking which of the 122 validation variables have been addressed, it is possible to identify both available and missing information needed to successfully develop or reuse an assay. The level of validation required for successful routine species monitoring will also differ on a case-by-case basis. For example, the *T. cristatus* assay was not placed on Level 5 of the validation scale (in part due to lack of reporting) but was still approved as an official survey method in the UK. As a general recommendation, authors should report as much information as possible on the conducted validation steps, either in the main text or in the supplementary material. Additionally, developers should consult existing guidelines for best practices along the validation workflow prior to assay design and fieldwork (Bustin et al., 2009; Goldberg et al., 2016; MacDonald & Sarre, 2017; Klymus et al., 2019). To make the validation process accessible, we provide a checklist in SI1. Furthermore, a website (https://edna-validation.com/) was created to summarise the cornerstones of the validation process and the validation scale. This website will function as a curated repository for newly developed assays and authors are encouraged to enter all 122 variables and the minimum criteria to rank assays and calculate their scoring percentages on the validation scale. The website will also serve as a living document when improvements in technology and/or our understanding of eDNA in the environment advance.

The 5-level validation scale designed here provides an overview of the capabilities and uncertainties of targeted eDNA assays. However, the binary data entry system cannot replace a close check of previous publications as it does not always allow a qualitative assessment. Details for validation variables are often spread across different sections in a publication or ambiguously displayed. Thus, the checklist can be used as standard reporting guidelines for targeted eDNA assays. It should be emphasised that for specific research questions and associated publications, minimal validation efforts may be sufficient. Nevertheless, thorough validation is needed to reduce uncertainties and overcome the limitations associated with eDNA-based species monitoring. Furthermore, it is important that practitioners consider how an assay can be modified (e.g. using different PCR reagents) and whether this changes its validation level.

The successful application of targeted eDNA assays for routine species detection and monitoring largely depends on the scientific community and the industry providing eDNA services. Laboratories participating in ring tests such as that proposed for metabarcoding (Blackman et al., 2019) can facilitate consensus on analysis standards. For now, assay developers must respond to queries and help troubleshoot reproducibility issues. Such engagement will facilitate the application of targeted eDNA assays by other users and outside their original geographic scope or academic context. Finally, it is necessary for both the scientific community and the commercial laboratories to communicate realistic applications and limitations to end-users, as often an assay is not bad *per se*, but simply unsuitable.

## Data availability

The literature database, the validation checklist compiled from it, the R code, and seven datasets derived from the checklist and used for analyses are available on Figshare at https://doi.org/10.6084/m9.figshare.12184860.v1 The literature database, the validation scale, and the full checklist are also available at https://edna-validation.com/.

## Acknowledgements

This article is based upon work from COST Action DNAqua-Net (CA15219), supported by the COST (European Cooperation in Science and Technology) programme, and RCB was also funded by SNF grant nr. 31003A_173074. We thank the workshop participants who convened on 26-27 March 2018 at the University of Innsbruck, Austria, where initial brainstorming for this manuscript occurred. We acknowledge additional guidance and input from Micaela Hellström, and are grateful to Cathryn Abbott for the improved design of Fig.1. We thank Nature Metrics Ltd. for support in hosting the independent website formed to operationalise the validation scale for future use.

## Conflict of interest

KB is the co-founder and CEO of Nature Metrics Ltd., a for profit company dedicated to the analysis of environmental DNA. HCR manages environmental DNA services for RSK ADAS Ltd., a for profit environmental consultancy. DS and MT are co-founders of Sinsoma GmbH., a for profit company dedicated to the analysis of environmental DNA.

## Author’s contributions

KB conceived the original idea, which was developed by all co-authors. BT compiled the literature database and BT, KD, LRH, HCR, RCB, DS, MT, and KB collected the data. BT analysed the data and led the writing of the manuscript for which first drafts of sections were provided by LH, RCB and KD. All authors contributed critically to the drafts and gave final approval for publication.

## Tables and Figures

**Box 1:**
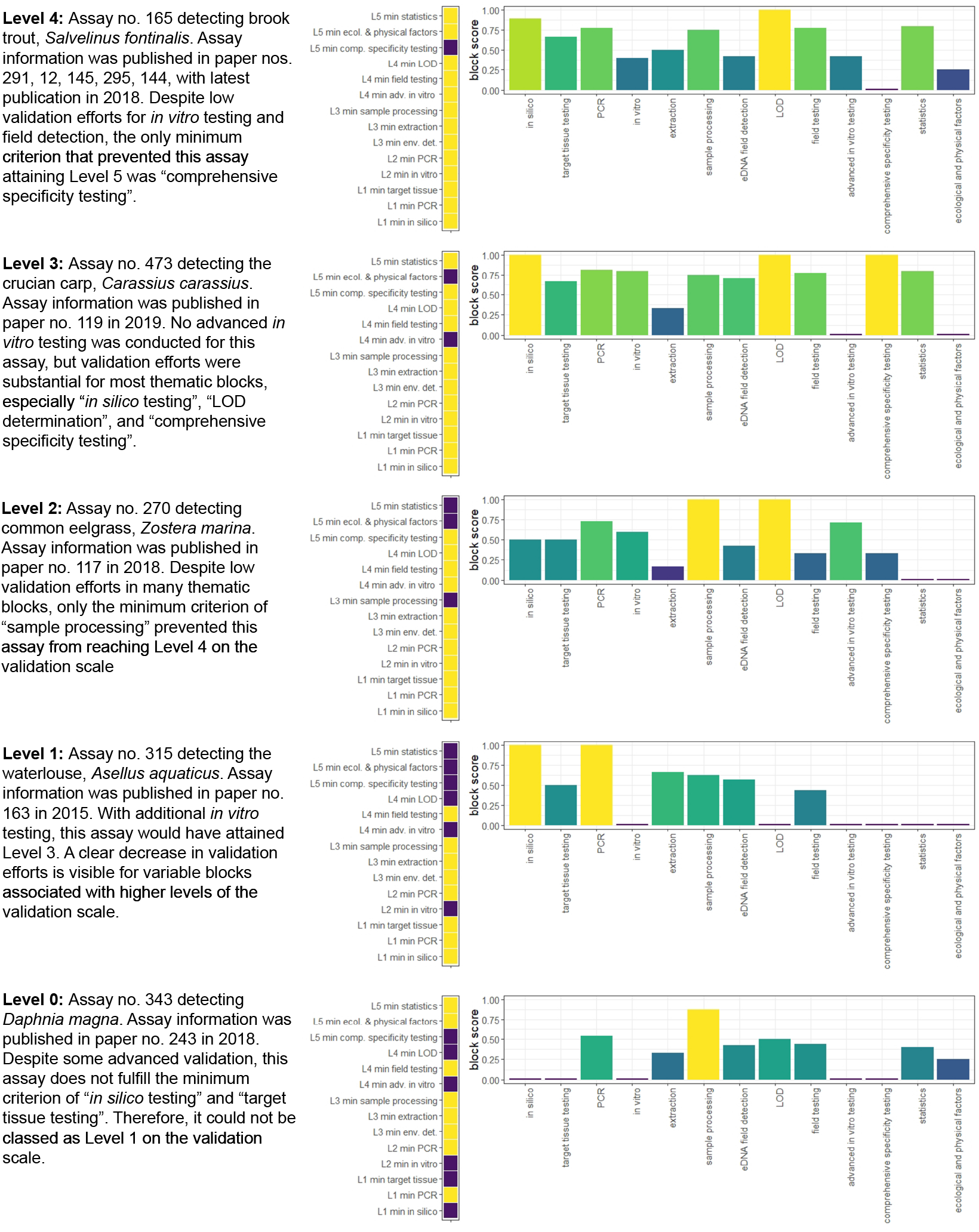
Examples of assays rated at Levels 0 to 4. The vertical tile plot shows which of the minimum criteria the assay fulfils (yellow tiles), and the bar chart gives the scoring percentage (i.e. the proportion of variables that were tested or reported) for each of the variable blocks. Bars are coloured according to the score obtained from a block with dark purple coding for “no validation” and yellow coding for “comprehensive validation”.

## Notes

https://doi.org/10.6084/m9.figshare.12184860.v1

https://edna-validation.com/

