## Supplementary material for "A validation scale to determine the readiness of environmental DNA assays for routine species monitoring": SI2

### Supporting Information 2

#### 2A) Compilation of the Literature Database

We compiled a literature database of published single-species eDNA assays, excluding work on microbial eDNA, metabarcoding and dietary studies. First, every paper listed on the "Environmental DNA: Tools and Resources" homepage of Washington State University (https://labs.wsu.edu/edna/edna-assays) as of 10 April 2019 was checked for suitability and added to the literature database. Afterwards, a literature search on Web of Science (http://apps.webofknowledge.com/) was conducted on 11 April 2019 using the following search criteria:

TS=("environmental DNA" OR "eDNA") NOT TS=("biofilm" OR "biofilms" OR "microbial" OR "bacterial" OR "microorganism" OR "microorganisms" OR "metabarcoding" OR "metagenomics" OR "next generation sequencing"); Databases= WOS, KJD, MEDLINE, RSCI, SCIELO; Timespan=2008-2019; Search language=English; Research Domain: science and technology; Document type: article, correction, book.

A list of 660 Web of Science entries was returned. Each of the publications was manually checked for suitability based on the exclusion criteria above, resulting in a set of 319 papers. During the assay validation process, it was necessary to include additional papers that originally published a primer set or that contained other information vital to the assay validation process. The final literature database contained 327 papers (https://doi.org/10.6084/m9.figshare.12184860.v1).

#### 2B) R packages used for analysis

The following R packages were used for data analysis: "ggplot2" (Wickham, 2016), "RColorBrewer"(Neuwirth, 2014), "scales" (Wickham & Seidel, 2019), "viridis" (Garnier, 2018), "rpart" (Therneau & Atkinson, 2019), "rpart.plot" (Milborrow, 2019), and "gridExtra" (Auguie, 2017), "dplyr" (Wickham, François, Henry, & Müller, 2019) and "hrbrthemes" (Rudis, 2019).
