## Supplementary material for "A validation scale to determine the readiness of environmental DNA assays for routine species monitoring": SI3

### Supporting Information 3

### Visualisation of descriptive variables included in the meta-analysis

Based on the data obtained for 546 assays from 327 published papers (https://doi.org/10.6084/m9.figshare.12184860.v1), the following figures summarise taxonomic diversity (Fig. SI3a), the number of papers containing validation information for a single assay (Fig. SI3b), the PCR type used for amplification (Fig. SI3c), the sample type from which eDNA was extracted (Fig. SI3d), and the target genes used for primer design and amplification (Fig. SI3e).

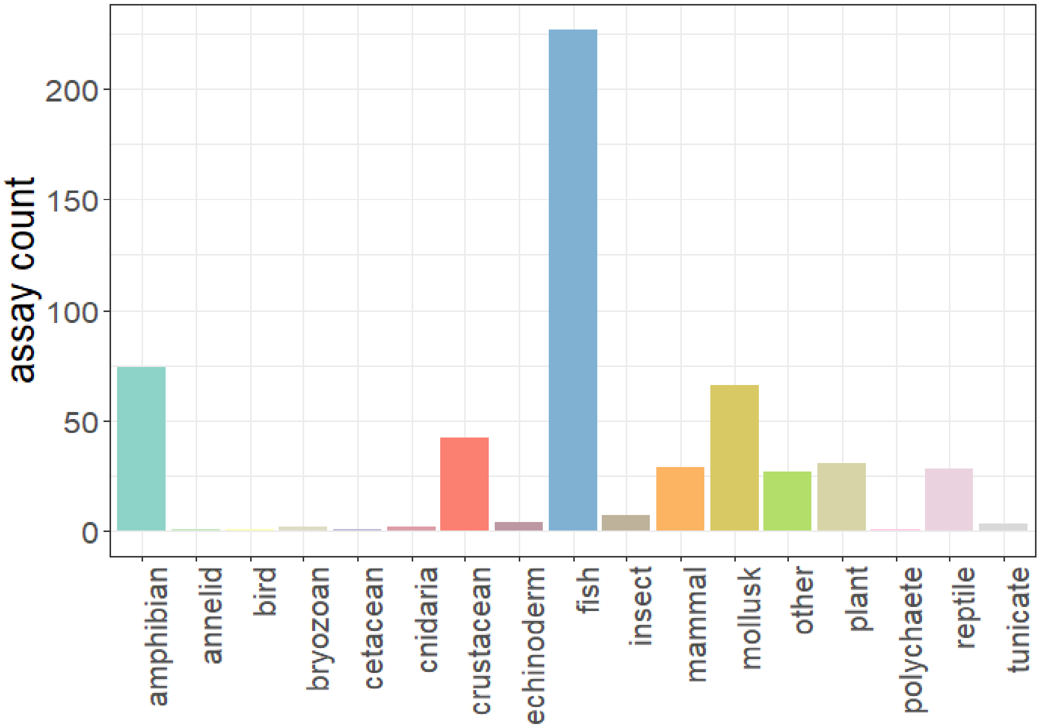


**Figure SI3a.** The number of assays developed per broad taxonomic group for the analysed dataset of 546 assays. Most assays were developed for fish species, followed by amphibians and molluscs.

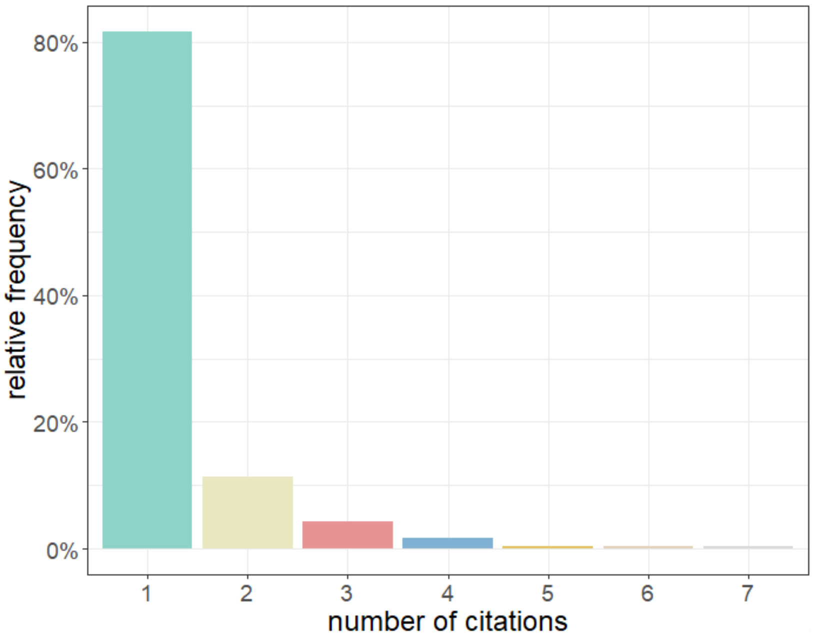


**Figure SI3b.** The number of papers containing information for a single assay in the analysed dataset of 546 assays. Over 80% of the examined assays occurred in only a single publication, but some were re-used, re-evaluated and developed further in up to seven papers.

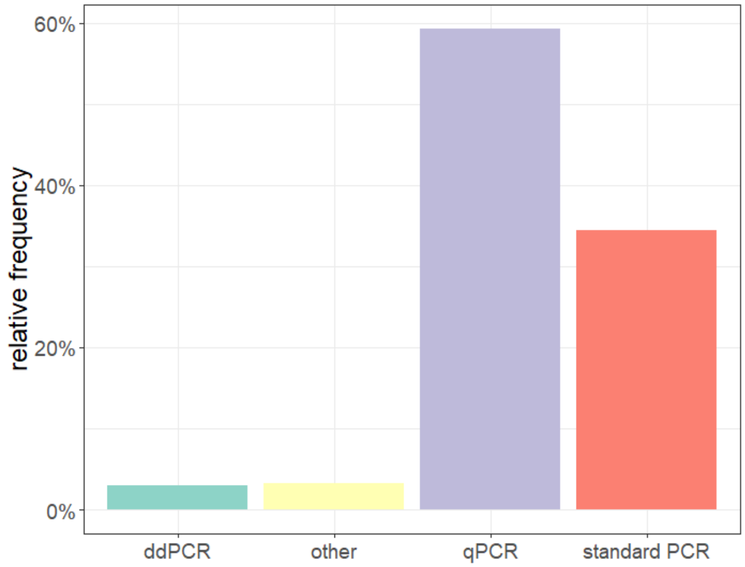


**Figure SI3c.** The PCR platform used for targeted eDNA amplification for all assays in the analysed dataset. Most assays were run on qPCR platforms.

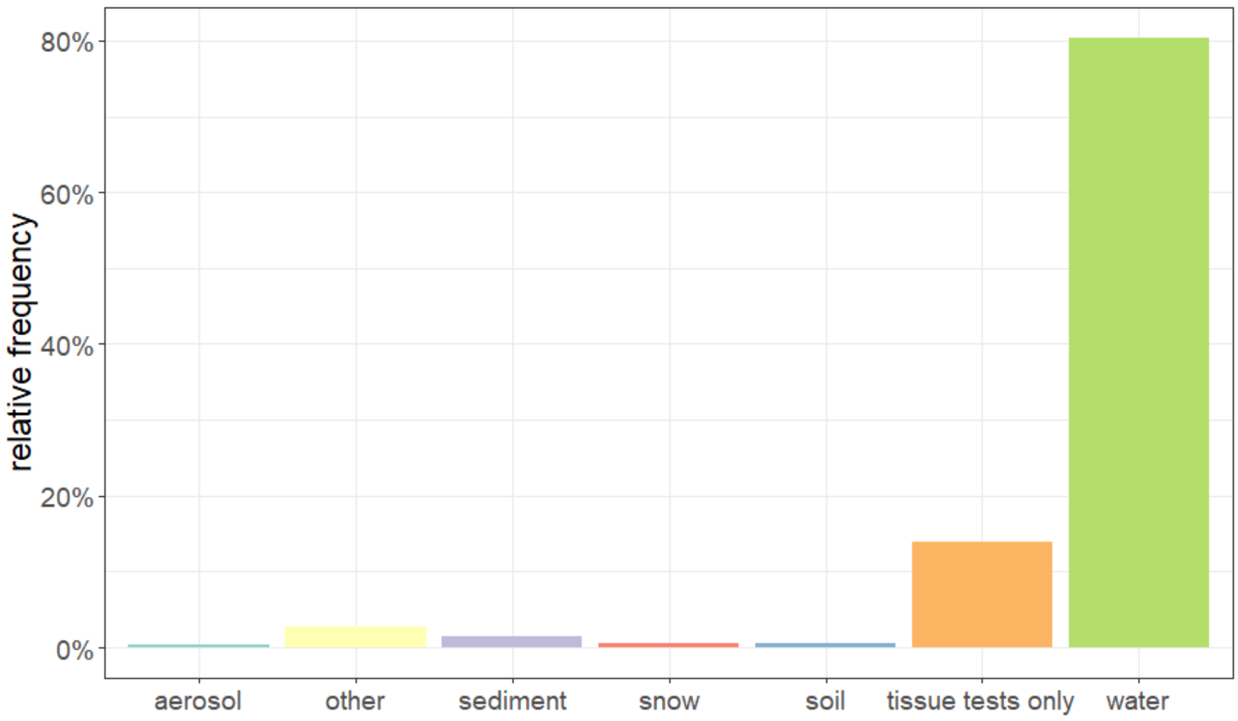


**Figure SI3d.** The sample type from which eDNA was extracted for the analysed dataset of 546 assays. In most cases, eDNA was extracted from water samples, and a considerable number of assays were only tested on tissue.

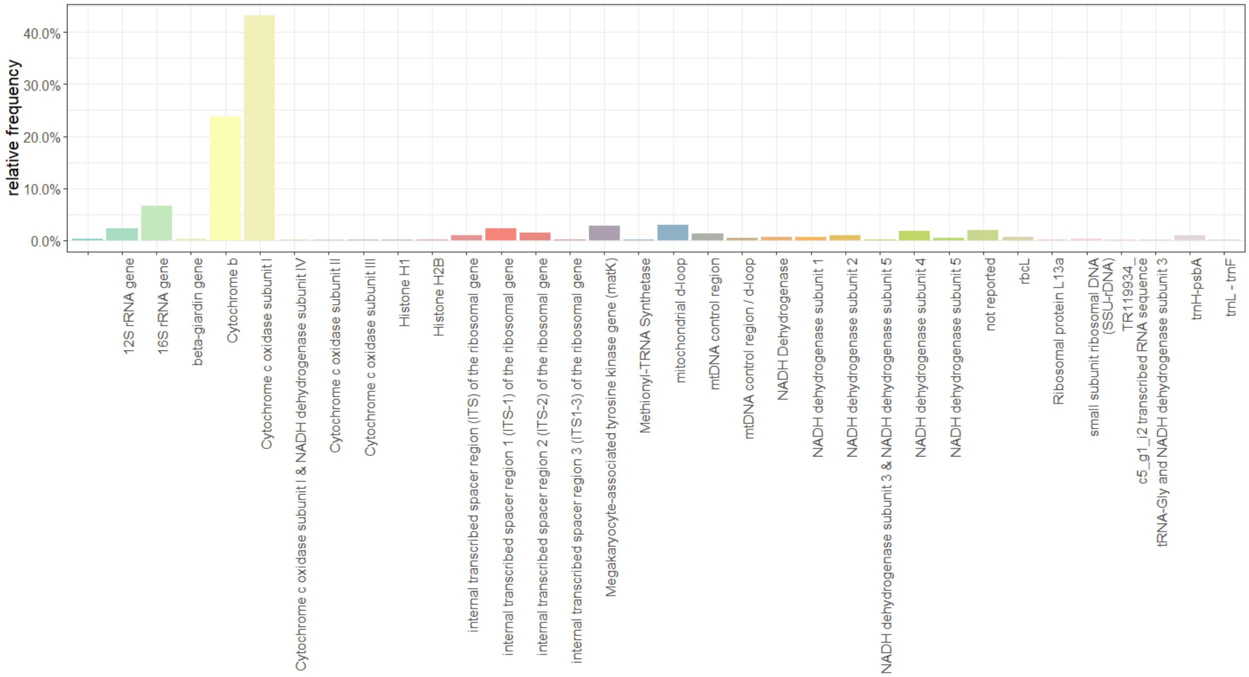


**Figure SI3e.** The target gene used for primer design and amplification with the 546 analysed assays. Most primers were designed to amplify fragments of the cytochrome c oxidase subunit I (*COI*) or the cytochrome b (*cytb*) genes. The first bar on the left denotes assays for which the target gene was not reported.
