## Supplementary material for "A validation scale to determine the readiness of environmental DNA assays for routine species monitoring": SI4

### Supporting Information 4

### Subjective assay rating versus objective assay rating

Each of the assays (https://doi.org/10.6084/m9.figshare.12184860.v1) was scored intuitively by a reviewer at the end of the evaluation process based on the basic validation scale version depicted in Figure 1. The scoring was possible from Level 1 to Level 5. To investigate the differences between subjective assay rating and objective assay rating based on the minimum criteria, the below figure (Figure SI4) was generated using R (R Core Team, 2019) and the package “viridis” (Garnier, 2018). The majority of assays classed as Level 2 or 3 by objective assay rating received the same subjective assay rating. However, most assays objectively classed as Level 1 were rated one or two levels higher by their subjective reviewers. In contrast, many assays objectively classed as Level 4 were subjectively rated as Level 3. Finally, two assays attained Level 5 based on subjective assay rating but were only classed as Level 3 or 4 objectively.

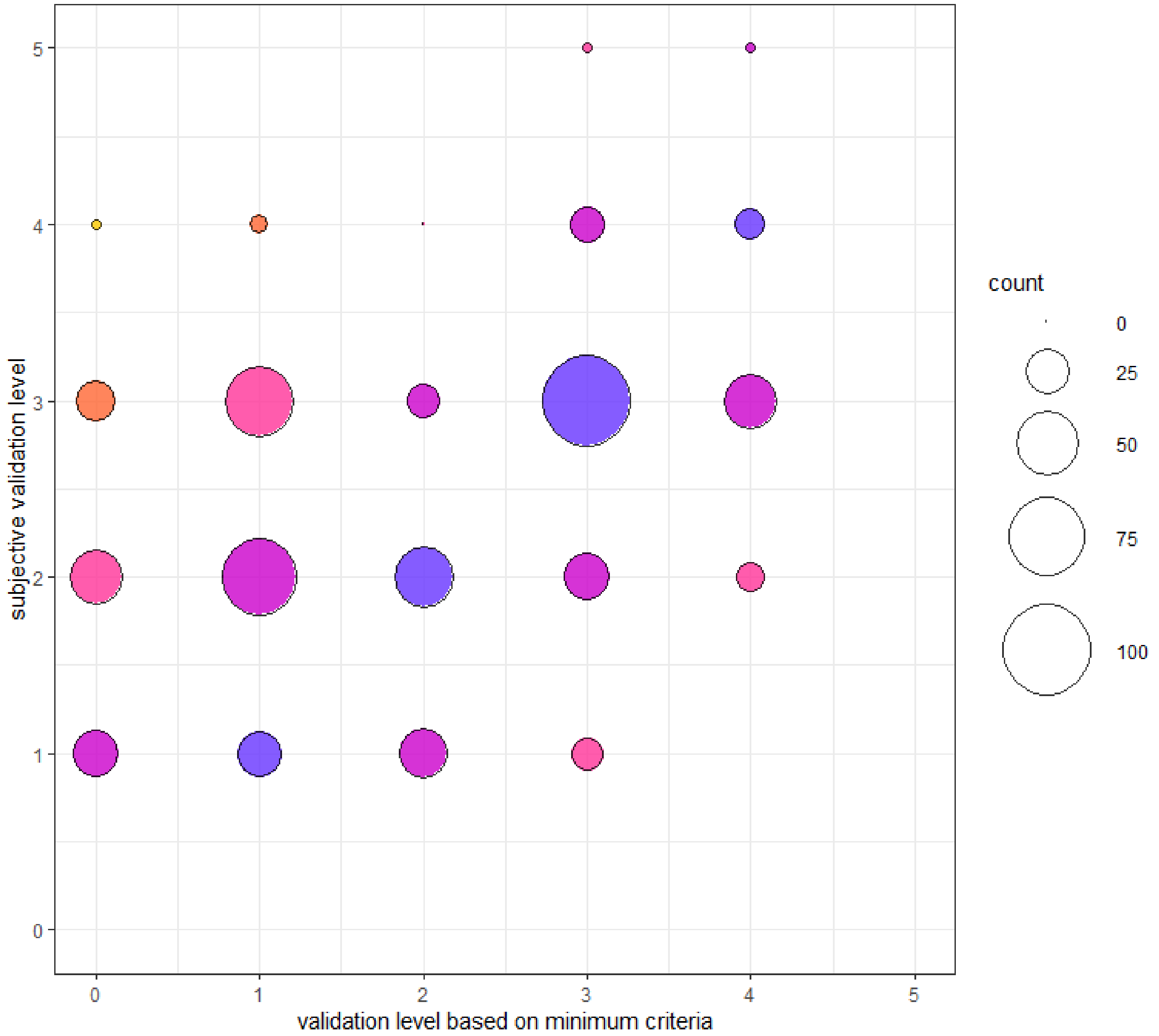


**Figure SI4:** Bubble plot comparing subjective and objective assay ratings. On the y-axis, the subjective assay rating is depicted (Levels 1 to 5 possible). On the x-axis the objective assay rating derived from the minimum validation criteria is depicted (Level 0 codes for assays that did not reach Level 1). The size of the bubble indicates the frequency of observed combinations of subjective and objective assay ratings. Bubble colour represents the difference in levels between subjective and objective assay ratings: purple (no difference), hot pink (one level), pink (two levels), orange (three levels), yellow (four levels).
