## Supplementary material for "A validation scale to determine the readiness of environmental DNA assays for routine species monitoring": SI5

### Supporting Information 5

### Classification Tree Analysis

Using the five levels of the validation scale as a response variable, a classification tree analysis was carried out (De'Ath & Fabricius, 2000). As the individual use of the 104 binary variables comprising the validation scale led to hardly comprehensible results (data not shown), scores of the checklist variables entered the analysis in a condensed form. For each of the 14 thematic variable blocks, an individual scoring percentage was calculated per assay: the sum of positive scores (yes/tested/reported) in a block was divided by the total number of applicable variables in the respective block. Hence, 14 block scoring percentages, i.e. values between 0 and 1, were generated for each of the 546 assays (https://doi.org/10.6084/m9.figshare.12184860.v1). These 14 block scoring percentages for each assay were used as explanatory variables for calculating a classification tree using R (R Core Team, 2019) and the packages "rpart" (Therneau & Atkinson, 2019) and "rpart.plot" (Milborrow, 2019).
